# Charting the cross-functional map between transcription factors and cancer metabolism

**DOI:** 10.1101/250399

**Authors:** Karin Ortmayr, Sébastien Dubuis, Mattia Zampieri

## Abstract

Transcriptional reprogramming of cellular metabolism is a hallmark feature of cancer. However, a systematic approach to study the role of transcription factors (TFs) in mediating cancer metabolic rewiring is missing. Here, we chart a genome-scale map of TF-metabolite associations in human using a new combined computational-experimental framework for large-scale metabolic profiling of adherent cell lines, and the integration of newly generated intracellular metabolic profiles of 54 cancer cell lines with transcriptomic and proteomic data. We unravel a large space of dependencies between TFs and central metabolic pathways, suggesting that the regulation of carbon metabolism in tumors may be more diverse and flexible than previously appreciated. This map provides an unprecedented resource to predict TFs responsible for metabolic transformation in patient-derived tumor samples, opening new opportunities in designing modulators of oncogenic TFs and in understanding disease etiology.

## Introduction

Transcription factors (TFs) are at the interface between the cell’s ability to sense and respond to external stimuli or changes in internal cell-state^1^. In cancer^2^ as well as other human diseases^3^, alterations in the activity of TFs can remodel the cellular signaling landscape and trigger metabolic reprogramming^4^ to meet the requirements for fast cell proliferation and cell transformation^5,6^. However, evidence linking alterations of cancer metabolism to TF dysfunction is often based on TF-binding sites upstream of metabolic enzymes^4^, without reporting on functional consequences of detected interactions. Mass-spectrometry based metabolomics approaches are powerful tools for the direct profiling of cell metabolism and to uncover mechanisms of transcriptional (in)activation of metabolic pathways^7–10^. Because of the limitations imposed by commonly used workflows, such as coverage, scalability and comparability, simultaneously quantifying the activity of TFs and metabolic pathways at genome- and large-scale remains a major challenge. Here, we develop a unique experimental workflow for the parallel profiling of the relative abundance of more than 2000 putatively annotated metabolites in morphologically diverse adherent mammalian cells. This approach overcomes several of the major limitations in generating large-scale comparative metabolic profiles across cell lines from different tissue types or in different conditions, and was applied here to profile 54 adherent cell lines from the NCI-60 panel^11^. To understand the origin of variance in the metabolome among different cancer cell types, we implemented a robust and scalable computational framework that integrates metabolomics profiles with previously published transcriptomic and proteomic datasets to resolve the flow of signaling information across multiple regulatory layers in the cell. This computational framework enables (i) systematically exploring regulation of metabolic pathways by TFs, (ii) reverse-engineering TF activity from *in vivo* metabolome profiles and (iii) predicting post-translational regulatory interactions between metabolites and TFs. Beyond contributing to the understanding of tumor-induced metabolic changes, this new systematic platform for charting genome-scale functional associations of metabolites with TFs in humans can introduce a new paradigm in the analysis of patient-derived metabolic profiles and the development of alternative strategies to counteract upstream reprogramming of cellular metabolism.

## Results

### Large-scale metabolic profiling of cancer cells reveals the interplay between transcriptional and metabolic heterogeneity

Tumor cells, in spite of similar genetic background or tissue of origin, can exhibit profoundly diverse metabolic phenotypes^12–14^. To systematically investigate the origin of heterogeneity in tumor metabolism we exploit the naturally occurring variability among a diverse set of cancer cell lines from the NCI-60 panel of tumor-derived cell lines^11^, representative of eight different tissue types. In spite of significant advancements in the rapid generation of high-resolution spectral profiles of cellular samples^15,16^, the accurate comparative profiling of intracellular metabolite abundances across large panels of different cell types is still a major challenge. Here, we present an innovative and robust workflow to enable large-scale metabolic profiling in adherent mammalian cells alongside with a scalable computational framework to compare molecular signatures across cell types with large differences in morphology and size. In contrast to classical metabolomics techniques^17,18^ employing small-scale cultivation formats, laborious and time-consuming extraction procedures, and additional quantification of cell numbers/volume, we use a 96-well plate cultivation format, rapid *in situ* metabolite extraction, automated time-lapse microscopy and flow-injection time-of-flight mass spectrometry^15^ (FIA-TOFMS) for high-throughput profiling of cell extract samples (Figure 1a).

**Figure 1.**
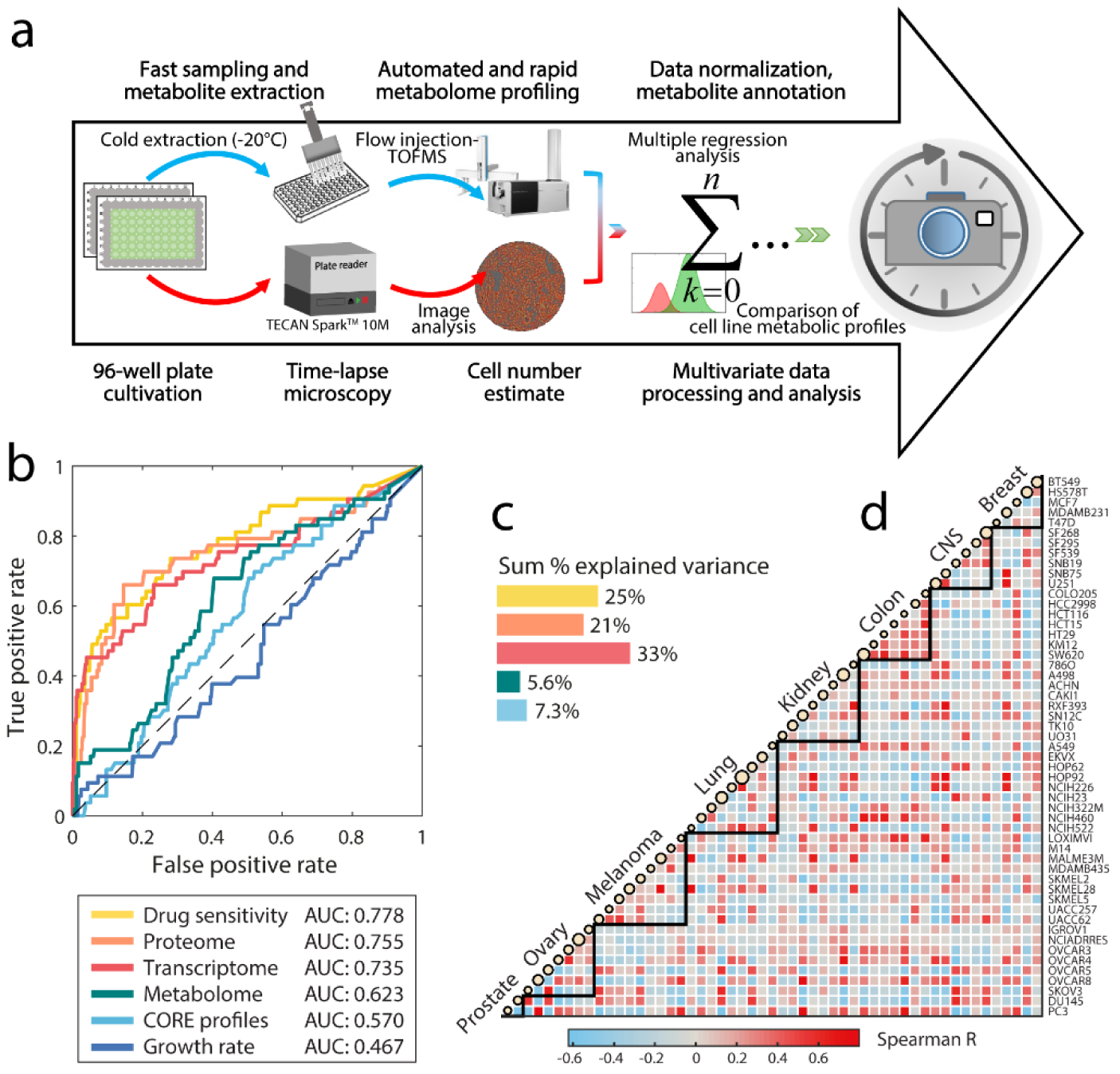
Comparative metabolome profiling in 54 adherent cancer cell lines. **(a)** Schematic overview of the combined workflow for high-throughput metabolome profiling in adherent cell lines. Multiple cell lines are cultivated in parallel to collect cell extracts for MS-based metabolome profiling (Supplementary Figure S1). A new software tool for the segmentation of bright-field microscopy images (Supplementary Note) is used for automated cell number quantification. Raw MS ion intensities are normalized using a multiple regression scheme (Supplementary Figure S2). **(b)** Signatures of tissue type in intracellular metabolome profiles compared with transcriptome, proteome, drug sensitivity, growth rate and the uptake and secretion of 140 metabolites (CORE profiles^14^). ROC curve analysis quantifies the performance of each of the data layers in predicting the tissue of origin (Supplementary Figure S3). **(c)** Major patterns in each data set were extracted by principal component analysis and tested individually for tissue signatures (Supplementary Figure S3). Colored bars indicate the sum of explained variance by tissue-associated principal components (AUC ≥ 0.75). **(d)** Pairwise similarity (Spearman correlation) of metabolome profiles among 53 cell lines from the NCI-60 panel. The size of circles on the diagonal is proportional to the growth rate (doubling time) of each cell line.

We optimized each step, from cultivation and extraction to MS analysis to be compatible with parallel 96-well processing. Different cell lines are seeded in triplicates at low cell density in 96-well microtiter plates, and are grown to confluence within 5 days (37°C, 5% CO_2_). Growth is continuously monitored by automated acquisition of bright-field microscopy images in 1.5-hour intervals. Using an in-house developed software for large-scale image analysis, for each cell line we can estimate average adherent cell size (i.e. the area covered on average by a single cell during adherent growth) and cell numbers at sampling time (see Supplementary Figure S1 and Supplementary Note). Replicate 96-well plates are sampled for metabolome analysis in 24-hour intervals. To facilitate sampling, increase the throughput and reduce the risk of sample processing artifacts, we collect metabolomics samples directly in the 96-well cultivation plate without any prior cell detachment (Supplementary Figure S1). Cell extracts were analyzed by FIA-TOFMS^15^, allowing rapid full-spectral acquisition within less than one minute per sample.

Normalization of large-scale non-targeted metabolomics profiles is a fundamental aspect of data analysis^19^, which becomes particularly challenging when comparing mammalian cell lines with large differences in cell size. We propose a two-fold approach: (i) measure metabolome profiles from the same cell line at different cell densities, enabling a multiple regression approach to decouple cell line-specific metabolic signatures from differences in extracted cell amount, plate-to-plate variance and background noise (Supplementary Figure S2), (ii) image analysis of bright-field microscopy images to correct for differences in cell volume between cell lines (Supplementary Figures S1-2). Notably, by integrating MS readouts at multiple cell densities and time points during growth, the resulting estimates of relative metabolite abundances are invariant to incubation time and cell densities, enabling the direct comparison between cells of different types and/or in different conditions. A detailed description of the workflow and our systematic multivariate approach to normalize metabolome data is provided in the Online methods.

Here, we compared the intracellular metabolomes of 54 adherent cell lines from eight different tissue types in the NCI-60 cancer cell line panel. Consistent with large differences in physiology^14,20^ and growth rates (Supplementary Data Table S1), we observed highly heterogeneous metabolome profiles across cell lines from the same tissue type (Figure 1d). Differences in growth rate correlated with the intracellular abundance of several intermediate metabolites in amino acid biosynthesis (Spearman |R| > 0.35, Supplementary Figure S3), reflecting essential biochemical requirements for rapid cancer cell proliferation^14^. Differently from metabolome profiles, analysis of previously published transcriptomics^21^ and proteomics^22^ data sets on the 54 cell lines revealed a stronger molecular signature associating with the tissue of origin (Figure 1b-d, Supplementary Data Table S2). The two major components of metabolic variance across the 54 cell lines (38% explained variance) were enriched for metabolites in nucleotide- and fatty acid metabolism as well as several signaling pathways (q-value < 0.05, Supplementary Figure S3), and correlated (Spearman |R| > 0.37) with the abundance of transcripts in signal transduction pathways regulating cell proliferation, adaptation, cell adhesion and migration (e.g. MAPK, HIF-1, PI3K-Akt, AMPK, Supplementary Figure S3). Altogether, these observations suggest metabolite abundances as closer readouts of cell physiology than mRNA or protein profiles, whilst pointing to a marked interplay between transcriptional regulation and the cell’s metabolic state.

### Mapping TF activity to metabolic phenotypes

To study the flow of signaling information between transcriptome and metabolome, we sought to quantify the functional interplay between different transcriptional programs and metabolic phenotypes. TFs can directly regulate metabolic fluxes by modulating enzyme abundance, thereby changing maximum flux capacity, or can indirectly affect substrate availability of proximal metabolic reactions, which can in turn result in local changes of fluxes and/or distal changes of reaction kinetics via post-translational regulatory mechanisms^23,24^. To gain insight into the functional regulatory role of TFs in metabolism, we aimed to establish correlations between TF activity and metabolite abundances across 53 cell lines, using metabolite abundances as reporters of the impact of TF activity on metabolic reactions^13,25^ (see Supplementary Figure S4 and Supplementary Discussion).

Because TF activity is a complex function of post-transcriptional and post-translational mechanisms, monitoring TF gene expression or protein levels is an inadequate proxy of TF activity^26,27^ (Supplementary Figure S4). Instead, we derived a relative measure of TF activities by integrating previously published transcript abundance data^21^ for 53 cell lines with a genome-scale network of TF-target gene interactions in human^28^ using Network Component Analysis (NCA)^29^ (Figure 2a, Supplementary Figure S4, Supplementary Data Table S3). By systematically correlating the activity of 728 TFs to relative levels of individual metabolites across cell lines (Figure 2a), we generated a network of TF-metabolite associations (Supplementary Data Table S4, Supplementary Figure S4). While in human cells most of the known TF binding sites in enzyme-encoding genes map to disease-related signaling pathways, our TF-metabolite network unraveled a complementary large space of associations involving intermediates in central metabolic pathways (Figure 2b). To functionally characterize the role of individual TFs in regulating distinct metabolic pathways, we tested TF-metabolite associations for an overrepresentation of intermediates in KEGG metabolic pathways (Figure 2c, Supplementary Figure S5, Supplementary Data Table S4). We discovered a significant enrichment (q-value < 0.05) for 677 TFs (mean: 8, median: 7 pathways per TF). Because of the characteristic difference in carbon metabolism (e.g. Warburg effect) between cancer- and normal cells^30,31^, oncogenic TFs that associate with central metabolic pathways are of particular interest (Figure 2c). A set of 38 oncogenic TFs showed significant associations with glycolysis, pentose phosphate pathway, tricarboxylic acid (TCA) cycle and/or oxidative phosphorylation (q-value < 0.05, Supplementary Figure S5), potentially reflecting a large space of regulatory interactions that can mediate the adaptation to diverse micro-environmental conditions and nutrient availability. For example, several TFs in the NF-κB signaling pathway (RelA, NF-κB2 and Bcl-3) exhibit a significant association with the pentose phosphate pathway (q-value < 0.05, Figure 2c), possibly reflecting the role of NF-κB complex as a mediator of redox homeostasis^32^ (see Supplementary Discussion).

**Figure 2.**
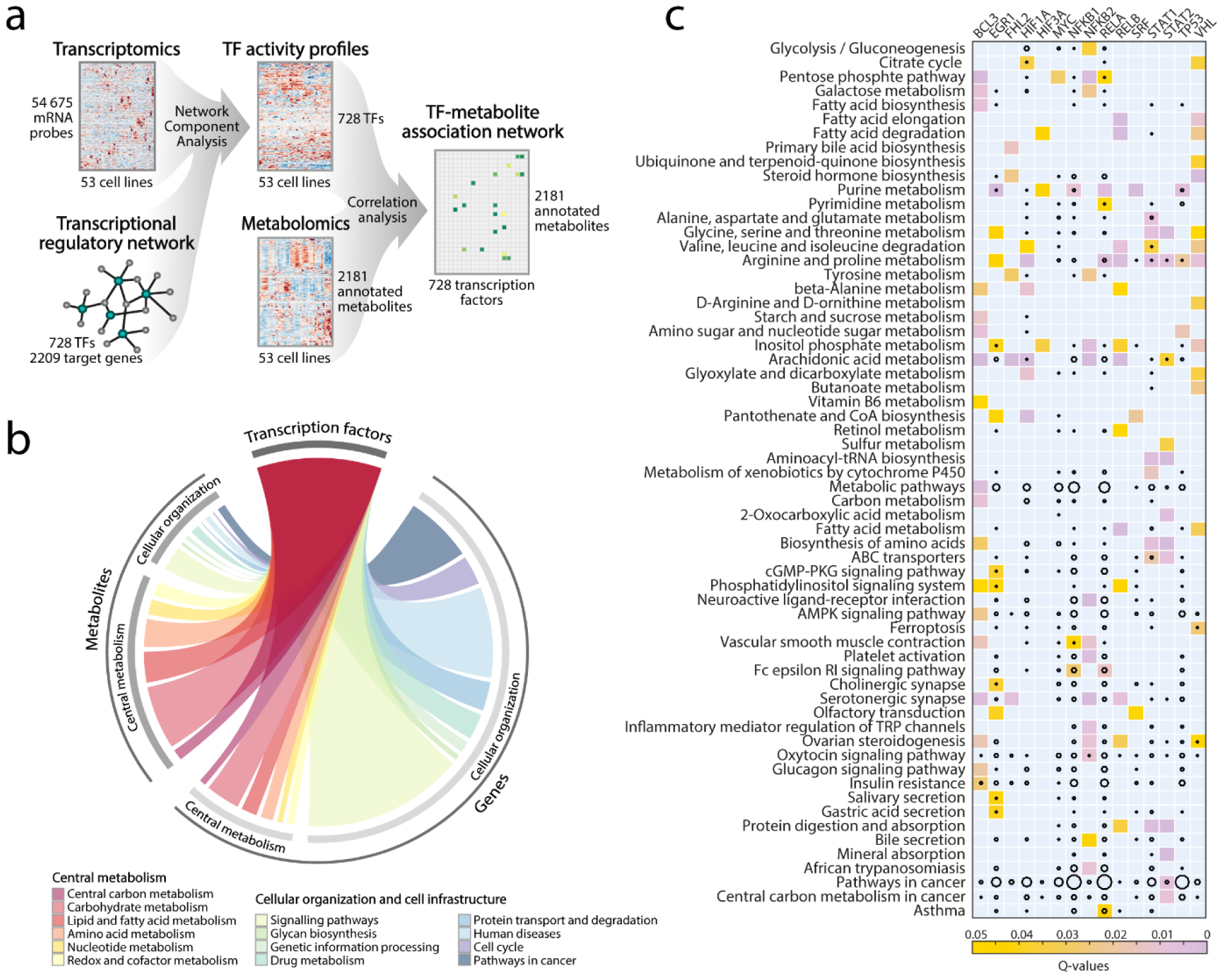
Inferring a TF-metabolite association network by integrating transcriptome and metabolome data. **(a)** Schematic overview of the computational framework for finding TF-metabolite associations. The activity of 728 human TFs was calculated using Network Component Analysis (NCA)^29^ which estimates TF activity from gene expression profiles of the NCI-60 cell lines and the transcriptional regulatory network between TFs and their target genes. In the second step, correlation analysis was used to find associations between TFs and metabolites (Figure 1). The resulting TF-metabolite association network resembles a scale-free network (Supplementary Figure S4). **(b)** Distribution of links to KEGG pathways in the TF-metabolite association network (left-hand section), and in the TF-gene regulatory network (right-hand section), respectively. KEGG pathways were grouped into super-pathways. Edge-size connecting the TF hub and a super-pathway reflects the number of links found in the network. **(c)** Metabolites associated with 15 known oncogenic TFs were tested for a significant overrepresentation in KEGG metabolic pathways (Supplementary Figure S5). Colored squares indicate the significance of the enrichment analysis (q-value < 0.05), while the size of black circles scales with the number of known TF-gene targets in the metabolic pathway.

The inferred associations between metabolic pathways and TFs are complementary to TF-gene regulatory networks. While the latter report on the regulatory capability of TFs to change enzyme abundance, coordinated changes between multiple metabolic intermediates and TFs can reveal the direct functional impact of TFs on pathway activity, and a TF’s potential in mediating the reprogramming of specific metabolic pathways in cancer cells. An emblematic example is tumor protein 53 (p53), one of the most well-studied tumor suppressor proteins^33,34^, for which significantly correlated metabolic intermediates in purine, arginine, proline- and nucleotide metabolism (q-value < 0.05) directly point to the functional role of the TF in sensing and allocating metabolic resources for cell division and DNA replication^33–35^. Even in cases where the role and target genes of TFs have been extensively characterized, the herein-proposed TF-metabolite associations can refine the condition-specific functional role of TFs in metabolism. For example, hypoxia-inducible factor 1 (HIF-1)^36^ is reported to act on regulatory elements upstream of nearly hundred enzymes in central metabolic pathways (Figure 2c). Here, we find a significant association with TCA cycle intermediates (q-value < 0.05), suggesting that changes in HIF-1 activity can directly transmit to TCA cycle. The predicted direct regulatory role is consistent with experimental evidence for a HIF-1-mediated activation of pyruvate dehydrogenase kinase (PDK) that actively represses TCA cycle activity^37,38^. In the following, we test the inferred TF-metabolite associations in an *in vivo* context, exploring the role of HIF-1 as a key oncodriver in clear-cell renal cell carcinoma.

### Inferring TFs mediating *in vivo* cancer metabolome rearrangements

Here, we ask whether the map of TF-metabolite associations found *in vitro* recapitulates metabolic rearrangements in an *in vivo* setting. To this end, we developed a computational framework that searches for TFs potentially responsible for metabolic differences between healthy and tumor tissue. The approach is based on a scoring function that evaluates the dot product between the TF-metabolite correlation matrix derived *in vitro*, and *in vivo* metabolite fold-changes between normal and cancer tissues (Supplementary Figure S6). Here, we applied this approach to analyze differences in metabolite abundances between clear-cell renal cell carcinoma (ccRCC) and proximal normal tissue samples in a cohort of 138 patients^7^. For each patient, we estimated the significance of a TF in explaining the observed metabolic changes, and ranked the 728 TFs according to the median of q-values across patients (Figure 3).

**Figure 3.**
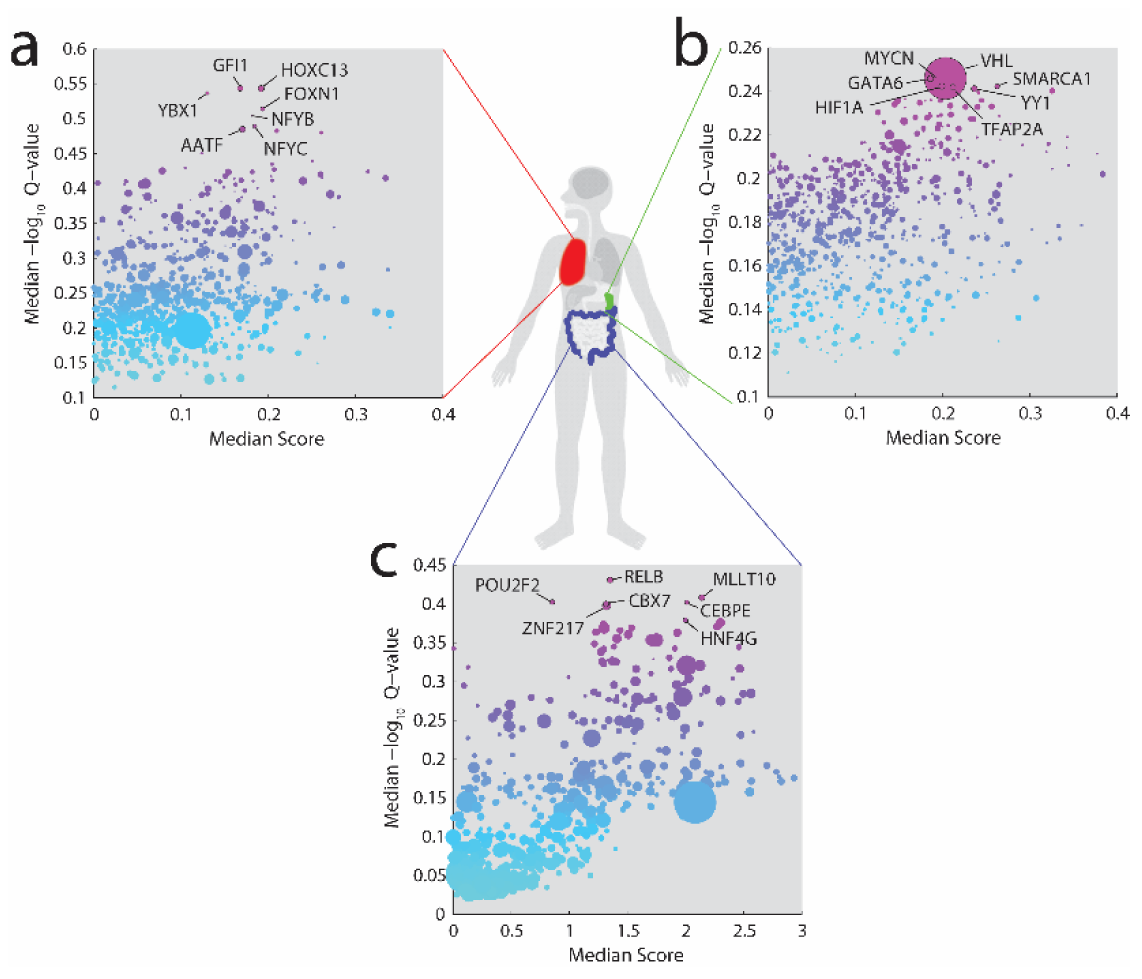
Inferring TF as mediators of *in vivo* metabolic changes. Prediction results in three previously published data sets comprising metabolic profiles of tumor vs. proximal healthy tissue from 21, 138 and 10 patients with lung cancer^43^ (panel **a**), clear-cell renal cell carcinoma^7^ (panel **b**), and colon cancer^13^ (panel **c**). Each dot represents the strength (x-axis, median score) and significance (y-axis, Q-value) of a TF in mediating *in vivo* metabolic changes. Dot-size indicates the frequency of mutations in the TF-encoding gene for the respective cancer type, derived from the Catalogue Of Somatic Mutations In Cancer (COSMIC)^78^. Gene-name labels are shown for the top 1% most significant hits (see Supplementary Discussion and Supplementary Figure S6).

Loss-of-function mutations in van Hippel Lindau (VHL) TF gene are the most frequent and specific genetic event observed in ccRCC^39^, entailing a hyper-activation of hypoxia-inducible factors (HIF-1, HIF-2 and HIF-3)^40^. In agreement with the genetic basis of ccRCC, we identified VHL and HIF-1α among the top 1% of TFs that potentially mediate metabolic rearrangement in ccRCC (Figure 3). Other top-ranking TFs include YY1 that can further stabilize HIF levels in VHL-null renal carcinoma^41^, and TFAP2A, whose gene is frequently hyper-methylated in ccRCC^42^ (see also Supplementary Discussion and Supplementary Figure S6). These results support the relevance of the previously inferred *in vitro* map of TF-metabolite associations and its potential clinical value in the interpretation of *in vivo* metabolome profiles of tumor tissue samples.

To further illustrate the potential of TF-metabolite associations in aiding the interpretation of tumor specific metabolic changes, we collected data from two additional studies monitoring tumor metabolic reprogramming in cohorts of 10 and 21 patients with colon^13^ and lung cancer^43^, respectively. In contrast to ccRCC, the most recurrent genetic events in colon and lung cancers are mutations in p53^44,45^, shared across many different tumor types. In such cases, our approach can dissect specific TFs, possibly effectors up- or downstream of p53, that can more directly explain tumor-specific metabolic changes. In colon cancer, we predicted an involvement of Mllt10, a downstream effector of Wnt signaling that is activated in many colorectal cancers^46^, as well as RelB, an NF-κB subunit which contributes to the rewiring of cancer metabolism through crosstalk with p53 and is involved in colitis-associated cancers^47^ (Figure 3 and Supplementary Discussion). Similarly, the metabolic changes observed in the lung cancer cohort highlight AATF, an upstream regulator of p53^48^, as well as NF-Y, a putative regulator of cancer metabolism that can act in complex with gain-of-function mutant p53 to increase DNA synthesis^49^ (Figure 3 and Supplementary Discussion). Taken together, *in vitro* associations between TFs and metabolites not only provide a means to functionally characterize the role of TFs across diverse cancer types, but could also complement genomic information and guide the analysis, molecular classification and interpretation of metabolome profiles from large patient cohorts.

### Systematic mapping of TF activity modulators

While we have shown that metabolic rearrangements in cancer can be described by changes in TF activity, the origin of such changes often remains elusive. Mutations in genes encoding TFs can be directly responsible for altered TF functionality, and in some cases even explain disease etiology.

However, the activation of new transcriptional programs is often an indirect response to changes in the abundance of internal effectors of cell signaling^50,51^. *In silico* models have proven extremely powerful in finding new allosteric interactions that can regulate enzyme activity ^52^ and in testing their *in vivo* functionality ^53^, but little progress has been made in the systematic mapping of effectors of TF activity. Here, we integrated three layers of biological information to obtain first insights into how TFs could be embedded in a regulatory network allowing the cell to activate distinct transcriptional programs in response to changes in the abundance of specific intracellular metabolites.

Besides mutations in TF-encoding genes, interactions with other proteins (co-factors or kinases/phosphates) and metabolites (e.g. allosteric binding) within the cell can repress or enhance TF activity. Despite the increased interest in metabolites as signaling molecules and their potential role in driving cellular transformation^50,54–56^, resolving the influence of metabolites on TF activity has ^57^. Here, we established an *in silico* framework for generating hypotheses on regulatory interactions between TFs, metabolites and kinases (Figure 4a). To that end, we used model-based fitting analysis to integrate TF activity and metabolome profiles with proteome data^22^ measuring the abundance of 100 TFs and 64 kinases/phosphatases across 53 cell lines. For each TF, we applied nonlinear regression analysis to determine whether variation in TF activity across the 53 cell lines could be modeled as a function of TF protein abundance and the activating or inhibiting action of individual metabolites and/or kinases. In total we tested 6,753,600 models, and found 1,888 interactions which significantly (FDR <= 0.1%) improved the explained variance in the activity of 96 TFs. Most of the inferred regulatory interactions (93%) involved the combined action of a kinase and a metabolite, while only few metabolites or kinases could alone significantly improve model-fitting (Figure 4b, Supplementary Data Table S5). This observation suggests that multiple coordinated regulatory mechanisms underlie the post-transcriptional regulation of TF activity. On average, each metabolite engaged in 8 interactions (median: 3), while kinases on the other hand were predicted to be more ubiquitous effectors (13-125 interactions per kinase) (Supplementary Figure S7).

**Figure 4.**
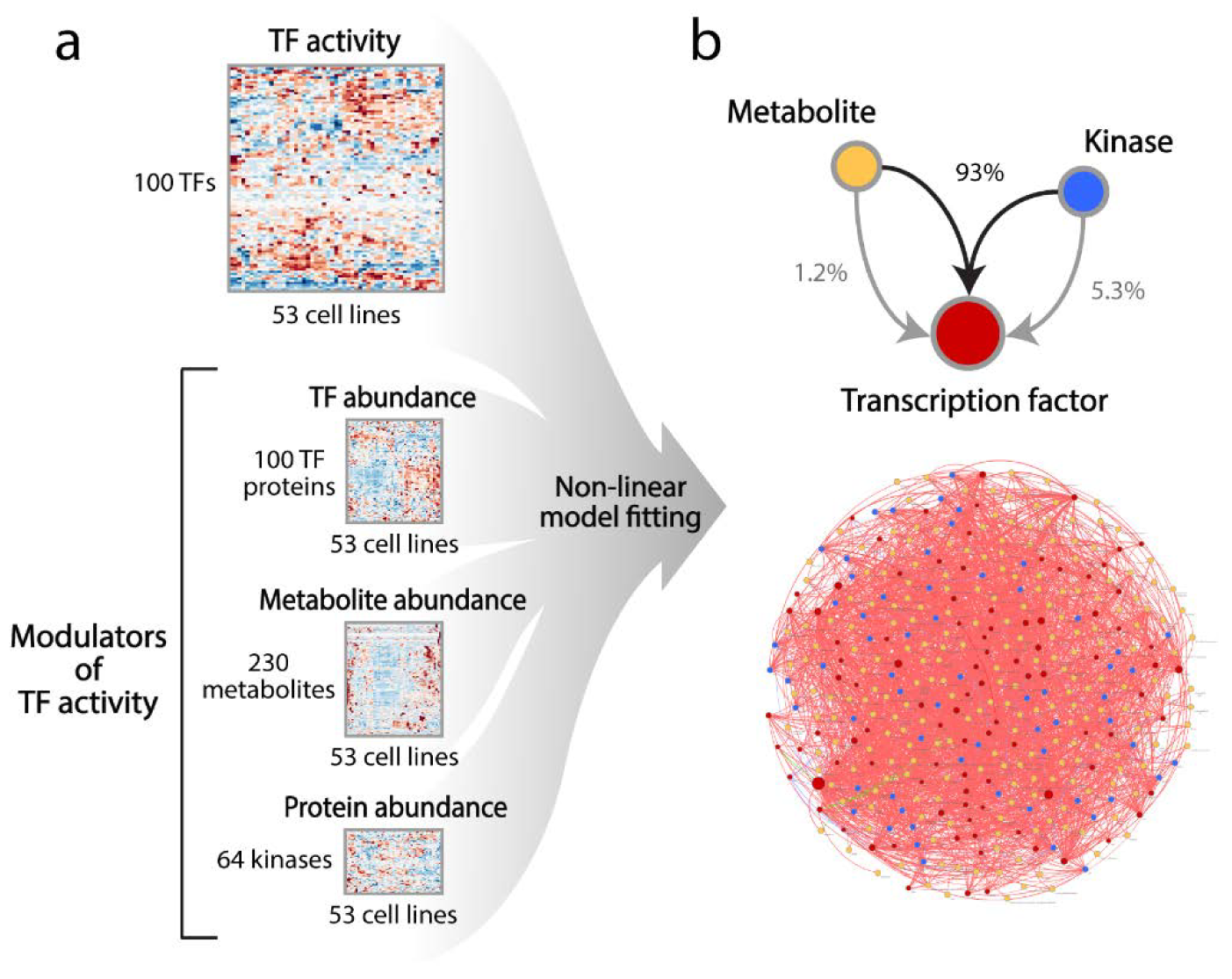
Modeling TF activity using an ensemble of non-linear models. **(a)** Schematic overview of the computational framework. Three layers of biological information are integrated using a model-based fitting approach: protein abundance of 100 TFs and 64 kinases, TFs activity derived from transcriptome data and relative levels of 230 metabolites across the 53 cell lines (see Online methods). **(b)** Predicted network of modulatory interactions between TFs (red nodes), metabolites (yellow nodes) and kinases (blue nodes). Most interactions involve the combined action of a metabolite and a kinase (93% of interactions). In 22 cases, changes in metabolite abundance alone were sufficient to improve the description of changes in TF activity (Supplementary Data Table S5).

This blueprint interaction map can serve as a new resource for generating hypotheses and designing target-oriented experimental approaches for in-depth mechanistic studies identifying metabolic regulators of transcriptional reprogramming. The most highly connected metabolite is choline (118 interactions with 36 TFs), substrate of the first step in the biosynthesis of phosphatidylcholine, a key component of eukaryotic cellular membranes. Recent studies have revealed a deep interplay between oncogenic signaling and choline metabolism, such that altered choline levels have become a metabolic hallmark of malignant transformation^58^. Our results support this key role, suggesting that choline availability could be sensed and can trigger a wide range of transcriptional adaptive processes. Further examples of predicted regulatory interactions are provided in the Supplementary Discussion.

Overall, our approach opens the door for a systematic investigation of a previously largely unexplored^59^ interaction space between transcriptional regulators and signaling effectors in human cells. The herein-presented experimental workflow for large-scale comparison of metabolomes in different cancer types, and *in silico* modeling of regulatory actions of metabolites and/or kinases can become an invaluable methodology in finding unique effectors that can trigger changes in the cell’s transcriptional program.

## Discussion

In recent years, increasing efforts have been made to understand the signals driving metabolic changes in cancer^13^. Transcriptional regulation is at the basis of the decision-making process of a cell and its ability to allocate resources necessary for cell transformation and proliferation. Genome sequencing and transcriptome technologies have revealed an intricate network of transcription factor–target gene interactions in which TF mutations often associate to disease states and aberrant metabolic phenotypes^7,60,61^. However, it is important to emphasize that gene regulatory interactions between enzymes and TFs per se are not sufficient to functionally regulate the activity of metabolic pathways. Key to unambiguously resolving regulatory circuits at the intersection with metabolism are methods searching for coordinated behavior between the different levels. Here, we integrate biological information across three layers: the transcriptome, the proteome, and the metabolome of 53 phenotypically diverse cell lines from the NCI-60 panel of human tumor cell lines. Key to this approach was the development of a combined experimental and computational framework (Figure 1a) that overcomes important limitations in large-scale metabolome screenings, including (i) the limited throughput and laborious sample preparation of classical metabolomics approaches in mammalian cell cultures^17,18^, (ii) the lack of scalable methods to adequately normalize metabolomics data across morphologically diverse cell types, and (iii) the need for systematic data integration strategies.

Here, by integrating cross-sectional omics data from diverse tumor cell lines, we constructed a global network model across three layers of biological information, exploiting the naturally occurring diversity in an *in vitro* cell line system. By analyzing the coordinated changes in baseline transcriptome, proteome and metabolome with the aid of a gene-regulatory network and model-based fitting analysis, we investigated the information flow from TFs to metabolic pathways and vice versa. The herein-constructed network of TF-metabolite associations provides an unbiased metabolic characterization of TF functions in human, and a more systematic understanding of functional regulatory interactions that can mediate metabolic adaptation in cancer cells. For many TFs, we found new regulatory associations with central metabolic pathways, suggesting a large space of transcriptional solutions by which cells can fulfill the anabolic and catabolic requirements for rapid proliferation^62^ and adaptation to nutrient limitations^63,64^. Secondly, we constructed a map of functionally relevant modulatory interactions between transcription factors, cell metabolism and cell signaling (kinases). Most interactions included the combined action of a metabolite and a kinase, acting as allosteric effectors or post-translational modifiers of TFs. An extensive network of these two types of physical interactions has emerged in model organisms such as *E. coli* or Yeast ^24,65,66^, and based on our findings we expect a similar picture to hold true in humans^59^. However, the identity and condition-dependent relevance of such interactions is much harder to determine experimentally in human cells than in unicellular model organisms. As a result, discoveries of TF-metabolite interactions in human cells have been sporadic^67^. The analysis and predictions proposed in this study offer new opportunities for a more systematic and exploratory discovery of TF post-translational regulatory interactions. The experimental and computational components of our approach are scalable and should in the future be applied to include a direct readout of kinase activity (e.g. phosphoproteomics data), broader protein coverage, larger cohorts of cell lines and more diverse environmental conditions. The relevance of TF-activity modulators extends beyond cancer metabolism, to the general understanding of how fluctuating micro-environments trigger coordinated responses of signaling pathways and culminate in the adaptive rewiring of the cell’s functional landscape.

In light of the central regulatory role of TFs in cellular organization, targeting TFs is an extremely attractive way to counteract global gene expression changes that underlie cancer development^68–70^. Endogenous metabolites capable of modulating TF activity could become invaluable chemical scaffolds to design new therapeutic molecules targeting oncogenic TFs, with the potential to overcome shortcomings related to targeting kinase-mediated signaling cascades^70^. The herein proposed workflow for large-scale metabolome profiling is directly applicable to the study of dynamic metabolic responses to external stimuli, and can scale to larger cohorts that are now within reach of other molecular profiling platforms^71^. Altogether, our work also suggests that, while clearly far from typical *in vivo* conditions, *in vitro* cell line systems represent an invaluable discovery tool to investigate metabolic regulatory mechanisms that can still generalize to *in vivo* conditions and clinical settings. The experimental and computational framework proposed in this study is applicable to other systems or diseases, providing us with an unprecedented tool to investigate the origin of metabolic dysregulation in human diseases.

## Acknowledgments

We thank the National Cancer Institute (NCI) for providing the cancer cell lines. This work was supported by a Worldwide Cancer Research (WCR-15-1058) project funding to M.Z., K.O. was funded by the Austrian Science Fund (FWF): FWF P26603 and FWF W1224 Doctoral Program BioToP – Biomolecular Technology of Proteins. We thank Uwe Sauer and Nicola Zamboni for supporting this work and providing laboratory facilities, Dimitris Christodoulou, Maren Diether, Nicola Zamboni and Victor Chubukov for helpful feedback and discussions.

## Contributions

M.Z. designed the project. S.D. and K.O. performed the metabolome experiments. M.Z. and K.O. analysed the data. S.D. and M.Z. designed and implemented the image analysis framework. M.Z. and K.O. wrote the manuscript. All authors contributed to preparing the manuscript.

## Materials and Methods

### Cell cultivation

The NCI-60 cancer cell lines were obtained from the National Cancer Institute (NCI, Bethesda, MD, USA). After thawing, the adherent cell lines were expanded in cell culture flasks (Nunc T75, Thermo Scientific) at 37°C and 5% CO_2_ in RPMI-1640 (Biological Industries, cat.no. 01-101-1A) supplemented with 5% fetal bovine serum (FBS, Sigma Aldrich, F6178), 2 mM L-glutamine (Gibco, cat.no. 25030024), 2 g/L D-glucose (Sigma Aldrich, cat.no. G8644), and 100 U/mL penicillin/streptomycin (P/S, Gibco, cat.no. 15140122). After two passages, the cells were transferred to fresh medium where FBS was replaced by dialyzed FBS (dFBS, Sigma Aldrich, cat.no. F0392) with a reduced content of low molecular weight compounds, to improve accuracy of metabolite quantification. Cells were maintained in medium with dFBS throughout all experiments. The starting cell density for metabolomics experiments was determined for each cell line. To this end, cells were plated in triplicates at eight different starting cell densities and incubated at 37°C and 5% CO_2_ for 3 days. On the third day, the medium was changed in all wells by aspirating the spent medium using a multi-channel aspirator, washing once with phosphate-buffered saline solution (PBS, pH 7.4, Gibco, cat.no. 10010015 at 37°C) using a multi-channel dispensing pipet, and finally filling each well again with 150 μL of fresh medium. The plate was imaged to determine the confluence (see below) immediately before and after media change, and after 72 hours. The starting cell density for metabolomics experiments was then chosen to guarantee a minimum of 20-30% cell confluence before media change, and approx. 80% confluence after 72 hours.

### Cell imaging and image analysis

We monitor cell growth by measuring cell confluence (i.e. area of the well covered by cells in percentage) directly from 96-well plates using automated time-lapse microscopy imaging. Every 1.5 hours, bright-field microscopy images of each well were acquired using a TECAN Spark 10M plate reader. In addition, we developed an image analysis framework to segment cells and determine the characteristic cell size area (i.e. average surface area of single adherent cells) for each cell line (Supplementary Figure S1). To quantify the number of extracted cells, confluence was divided by the characteristic cell size (Supplementary Figure S1). A detailed description and validation of the algorithm used for estimating cell numbers from bright-field microscopy images is provided in the Supplementary Note, alongside with a Matlab code. Of note, this approach has several important advantages, in that it is non-destructive, and allows quantifying cell growth and cell numbers without any manual sample manipulation.

### Metabolomics experiments

Cells were plated in triplicates in 150 μL of RPMI1640 medium (5% dFBS, 2 g/L glucose, 2 mM glutamine, 1% P/S) in 96-microtiter well plates. After an initial growth phase, the medium in each well was renewed on the third day, and the cultures were subsequently monitored for four more days (96 hours). Ten replicate plates were prepared in each experiment to allow for generating metabolomics samples at five different time-points (immediately before media change, and at 24, 48, 72 and 96 hours after media change), and one additional plate for continuous growth monitoring (TECAN Spark 10M, 37°C, 5% CO_2_).

At each sampling time point, two replicate 96-well plates were processed (plates A and B). Plate A was used to generate cell extracts, by (1) removing the spent medium, (2) washing once with 75 mM ammonium carbonate (pH 7.4), and (3) adding ice-cold extraction solvent (40% methanol, 40% acetonitrile, 20% water, 25 μM phenyl hydrazine^72^). Finally, the plate is sealed, incubated at −20°C for one hour, and subsequently stored at −80°C until MS analysis. Plate B undergoes the same processing steps, except for the last one, where each well is filled with 37°C PBS (pH 7.4), and the plate is immediately imaged to determine cell confluence for subsequent normalization of MS spectra.

Immediately prior to MS analysis, the plates were thawed on ice, and the extracted cells were scraped off the bottom of each well using a multi-channel pipet with wide-bore tips. Next, the cell extracts were transferred to 96-well plates with conical bottom and centrifuged at 4°C, 4000 rpm for 5 min to separate cell debris. Finally, pooled cell extracts for each experiment (“master mixes”, pooled from five cell lines processed within the same experiment) as well as aliquots of unused extraction solvent were added to each measurement plate as control samples, and the plates were sealed and stored at 4°C until injection.

### Metabolome profiling using FIA-TOFMS

Cell extract samples were analyzed by flow-injection analysis time-of-flight mass spectrometry (FIA-TOFMS) on an Agilent 6550 iFunnel Q-TOF LC/MS System (Agilent Technologies, Santa Clara, CA, USA), as described by Fuhrer et al^15^. This method allows the generation of high-resolution spectral profiles in less than one minute per sample, allowing for sensitive high-throughput profiling of large sample collections. In brief, a defined sample volume of 5 μL is injected using a Gerstel MPS2 autosampler into a constant flow of isopropanol/water (60:40, v/v) buffered with 5 mM ammonium carbonate (pH 9), containing two compounds for online mass axis correction: 3-Amino-1-propanesulfonic acid, (138.0230374 m/z, Sigma Aldrich, cat. no. A76109) and hexakis(1H,1H,3H-tetrafluoropropoxy)phosphazine (940.0003763 m/z, HP-0921, Agilent Technologies, Santa Clara, CA, USA). The sample plug is delivered directly to the ion source for ionization in negative mode (325°C source temperature, 5 L/min drying gas, 30 psig nebulizer pressure, 175 V fragmentor voltage, 65 V skimmer voltage). Mass spectra were recorded in the mass range 50-1000 m/z in 4 GHz high-resolution mode with an acquisition rate of 1.4 spectra per second. Raw MS profiles were processed to align spectra and pick centroid ion masses using an in-house data processing environment in Matlab R2015b (The Mathworks, Natick).

### Metabolite annotation

Measured ions were putatively annotated by matching mass-to-charge ratios to a reference list of calculated masses of metabolites listed in the Human Metabolome Database (HMDB) and the genome-scale reconstruction of human metabolism^73^ (Recon2) within 0.003 amu mass accuracy. The reference mass list was generated from the respective sum formulae, considering deprotonation as the most prevalent mode of ionization in the chosen acquisition conditions. To allow for the annotation of α-keto acid derivatives formed in presence of phenyl hydrazine in the extraction solvent^72^, sum formulae for the phenylhydrazone derivatives (+C_6_H_8_N_2_-H_2_O) of a total of 30 α-keto acid compounds (selected via KEGG SimComp search http://www.genome.jp/tools/simcomp/) were added to the metabolite list for annotation. The final list of putatively annotated metabolites consisted of 689 and 5949 unique compound IDs from Recon2, and HMDB, respectively.

### Data normalization

We corrected for systematic errors using a two-step regression model to disentangle the contributions of extracted cell numbers, plate-to-plate variance, instrumental and background noise from the actual variance in metabolite abundances between cell types (Supplementary Figure S2). To this end, raw data were first corrected for instrument drift by normalizing for possible batch/plate effects. Each plate contains 12 pooled cell extract samples prepared from the 5 different cell lines in each experiment (i.e. batch). Measured intensities for each annotated ion are modeled as follows:

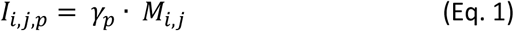

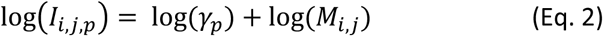

Where *I_i,j,p_* is the measured intensity for ion I, in pooled sample j and plate p, *γ_p_* is the scaling factor associated to each plate and *M_i,j_* represents the actual abundance of metabolite i in sample j. By using a linear regression scheme we can estimate both parameters (*γ_p_* and *M_i,j_*) within an unknown scaling factor. After correcting for possible instrumental artifacts, we implemented a second step in order to derive comparative measurements of metabolite abundance for each cell line. Here, we follow each cell line along the linear growth phase, sampling every 24 h across 5 days. We typically obtain 25 data points for each cell line at different cell densities. The expectation is that the signal measured for any ion of biological origin (i.e. a genuine metabolite) would increase linearly as the number of cells in the sample increases. The proportionality (i.e α parameter) between ion intensity and extracted cells depend on the intracellular concentration of the metabolite. Here, by implementing a multiple regression scheme, we estimate the relative abundance of a given metabolite in each of the cell lines, α (Supplementary Figure S2), alongside with its standard error. A linear regression model describes variation in ion intensity as a linear function of cells extracted (α) and a constant parameter (β) that capture MS background noise. For each ion, the α values are specific of each cell line while the constant term in the model is fixed (i.e. expected ion signal at zero confluence that is independent of the cell line).

Because of the large number of measurements for each cell line at different cell densities, we can apply a multiple regression analysis (*fitlm* function in Matlab) to infer all model parameters αs and β at once, by minimizing the Euclidian distance between measured metabolite intensities and model predictions. For each metabolite, we solve the following linear model, including all 54 cell lines:

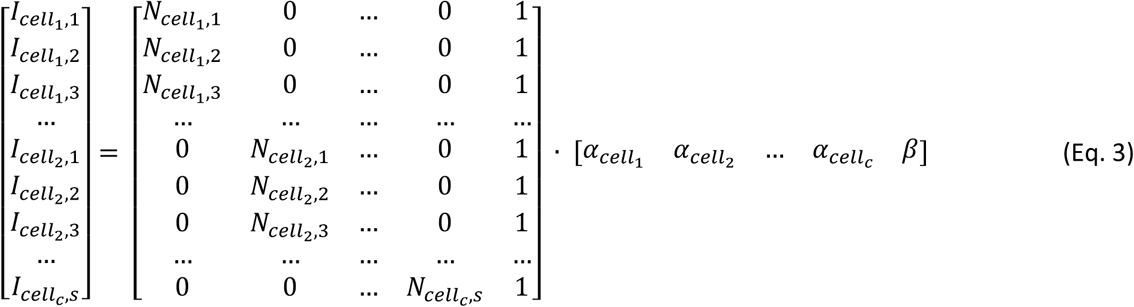

Where I_cell c,s_ is the measured metabolite intensity in sample s of cell line c, N_cell c,s_ is the number of cells extracted in sample s of cell line c, and αs (for each cell line) and β are the unknown parameters to be fitted. The number of cells per sample is derived from the confluence measurements at sampling, divided by the average cell size area determined using our image segmentation analysis (see Supplementary Figure S2 and Supplementary Note). After this step, we retained 2181 ions with a regression p-value below a threshold value of 3.4e-7 (adjusted by the number of cell lines and ions) in at least one cell line, and that showed a significant dependency with the extracted cell number in more than 80% of cell lines (Supplementary Figure S2). Of note, we found that prior to normalization, the variance across three biological replicates at the same time-point was equally low in cell confluence (median: 7.4% CV) and raw ion intensities (median: 13%, Supplementary Figure S2), reflecting the high quality of MS measurements.

In the third and last step, we take into account systematic changes in metabolite abundances related to differences in cell size (i.e. cell volume) between the 54 cell lines to derive comparative estimates of intracellular metabolite concentration. Principal component analysis of relative metabolite abundance per cell revealed a strong trend across the 54 cell lines (PC1, 58.9% explained variance, Supplementary Figure S2), which strongly correlates with cell line volumes derived from cell diameters measured in ^74^ as well as with the herein determined adherent cell size area (Supplementary Figure S2, see Supplementary Note). The transitive correlation between adherent cell size and the spherical cell volume in suspension indicates that adherent cell height can be approximated as a constant. To correct MS data for differences in cell line volumes, we selected 987 ions that showed a significant and strong correlation (Pearson’s r > 0.8, p < 0.05, Supplementary Figure S2) with PC1. KEGG pathway enrichment analysis showed that these putatively annotated metabolites were strongly overrepresented in fatty acid metabolism (Supplementary Figure S2), consistent with the expected linear dependency between cell membrane surface (i.e. phospholipid content) and cell volume of adherent cells. For each ion, cell line-specific α-values of the selected metabolites across all 54 cell lines were used to calculate a consensus correction factor for each cell line by taking the mean across the 987 ions. To apply the cell volume correction to the full data set, we divided the cell line-specific α-values for each ion by the consensus correction factor.

Finally, the corrected α-values were normalized using Z-score normalization:

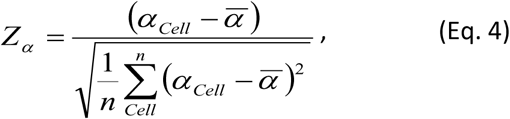

where n is the number of cell lines (i.e. 54).

The final normalized data set is provided in Supplementary Data Table S1 alongside with p-values and standard errors derived from regression analysis. Missing values (‘NaN’) correspond to cases where the measured ion abundance for the annotated metabolite was close to the background level in the cell line, and can be considered as zero for further analysis. In cell lines where a significant dependency (p < 3.4e-7) of given metabolite’s abundance with cell number could be robustly determined and exceeded the background noise, relative standard errors of α calculated during fitting analysis were below 20% (median: 11%, Supplementary Figure S2).

### Analysis of tissue signature in transcriptomics, proteomics, metabolomics and drug sensitivity data sets

Principal component analysis (PCA) was used to dissect each data set into major trends, and quantify the extent of variance explained by each of these principal components (Supplementary Figure S3). The PC scores obtained for the different cell lines in each principal component and data set were then analyzed for signatures of individual tissue types based on the pairwise Euclidean distance between cell lines. For each cell line, the enrichment of cell lines from the same tissue type among its nearest neighbors (i.e. smallest distance scores) was tested using a hypergeometric test. The resulting probabilities (p-values) from enrichment analysis were subjected to receiver-operator-characteristic (ROC) curve analysis to evaluate the performance of a given principal component as a predictor of the tissue type (Supplementary Table S2). A principal component was assigned a tissue signature when the area under the ROC curve (AUC) exceeded a value of 0.8 for any tissue type, and the fraction of explained variance was summed up to obtain the overall percentage of variance explained by tissue type in each data set (Figure 1c).

### Estimating TF activity using Network Component Analysis (NCA)

Originally established by Liao et al.^29^, NCA provides a mathematical framework for reconstructing TF regulatory signals (TF activity) from gene expression profiles. Here, we adopted sparseNCA implementation by Noor et al.^75^ (Matlab code downloaded from https://sites.google.com/site/aminanoor/softwares). This methodology adopts a mathematical model to approximate TF-target regulatory interactions and integrates prior network information with the expression of target genes across multiple conditions to regress the activity of the respective TFs, delivering a relative measure of TF activity. We obtained normalized gene expression profiles across the NCI-60 cell lines from Gene Expression Omnibus (accession number GSE32474), containing 54,675 mRNA probes. TRRUST database^28^ served as the source of TF-target gene interactions relevant in human, including 748 human TFs and 1,975 non-TF gene targets. Intersecting these two resources, we assembled a network of 2,209 unique genes corresponding to 5490 mRNA probes that match target genes of 728 TFs in the TRRUST database (Supplementary Figure S4). We implemented a bootstrapping approach to account for incompleteness of the regulatory network (i.e. missing regulatory interactions), and the fact that there may be multiple optima in the solution space. To this end, for each TF we randomly selected 48 additional TFs and constructed a sub-network containing the 49 TFs and their target genes. Because growth-rate has a pleiotropic effect on gene expression, here reflected in the correlation between first principal component of gene expression data and cell line growth rates (Supplementary Figure S4), we decouple TF activity from the confounding effect of growth-rate by adding an additional TF that targets all genes. This fictitious TF mimics the general effect of proliferation rates on transcription. As a result, each TF is embedded in a sub-network of 50 TFs and their target genes from the full network. Ten such sub-networks were created randomly for each TF to apply NCA. In this bootstrapping scheme, each TF was sampled in on average 490 subnetworks (permutations, min. 423, max. 556 data points per TF). In the final data set, we normalized the calculated TF activity to the maximum across all permutations, and finally calculated the median TF activity and its standard deviation for each TF and cell line (Supplementary Figure S4). It is worth noting that the estimates we obtain with this approach are correct within an unknown scaling factor, and hence we determine a relative measure of activity for each of the 728 TFs across the NCI-60 cell lines.

### TF-metabolite association network and inferring TF involvement from *in vivo* metabolic changes

In order to find metabolites whose relative abundances correlate with TF activity, we calculated pairwise Spearman correlations between all 2181 annotated metabolites and 728 TFs across the 54 cell lines. In order to control the false discovery rate (FDR) among network links, we used a bootstrapping approach to calculate 99.5 and 99.9% confidence intervals of correlation coefficients after randomizing the data set. To that end, the cell lines in the metabolome data set were randomized by resampling 100 times with replacement, and Spearman correlation coefficients were calculated for each randomized data set. Correlation coefficients yielding 99.5 and 99.9% confidence intervals (0.5 and 0.1% FDR, respectively) were obtained from the pooled list of absolute correlation coefficients by finding the smallest correlation coefficient that exceeds the maximum value among 99.5 and 99.9% lowest correlation coefficients, respectively).

Based on this association network, the procedure used to estimate the contribution of each of the 728 TFs in mediating metabolic changes observed *in vivo*, consists of 3 main steps: (i) For each patient log_2_ fold-changes of detected metabolites between cancer and adjacent normal tissue are estimated (FC^p^), (ii) The dot product between metabolite fold-changes and the TF-metabolite correlation vector (c_TF_, product of correlation R and −log_10_ p) estimated from *in vitro* cell lines is computed for each TF 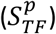, (iii) the significance is estimated using a permutation test, where metabolite order is shuffled 10.000 times, the dot product is estimated for each random permutation 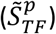 follows:

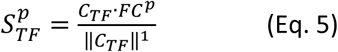

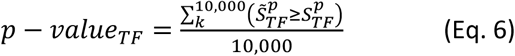

P-values are corrected for multiple tests by q-value estimation^76^, and the median across patients is calculated. Notably, when analyzing the data published in ^7^ we excluded all detected metabolites with more than 10 missing values across patient samples.

### KEGG enrichment analysis of TF-metabolite correlation signatures

To assess the over-representation of TF-metabolite associations in KEGG metabolic pathways, the pairwise correlation between TF activity and metabolite relative abundance across cell lines were rank-transformed. A statistical score that models the probability of a KEGG pathway to be significantly associated to a TF is based on the collective activities of multiple metabolites in a pathway following the approach described in ^77^. The significance of the rank distribution of all metabolites within the same KEGG pathway is tested by means of an iterative hypergeometric test, indicating the statistical significance of metabolic intermediates of a common metabolic pathway (e.g. TCA cycle) being distributed toward the top ranking ones. P-values were corrected for multiple tests by q-value estimation ^76^.

### Prediction of metabolite-TF effectors

Because of the poor correlation between protein levels and NCA estimated TF activities (Supplementary Figure S4), we systematically investigate whether variation in TF activities across cell lines could be explained by alternative models. Instead of assuming a base model where the TF activity is a simple linear function of protein abundance, we model TF activity as a function of TF protein levels and the post-translational regulatory functions of metabolites and/or kinases, individually or in combination as follows:

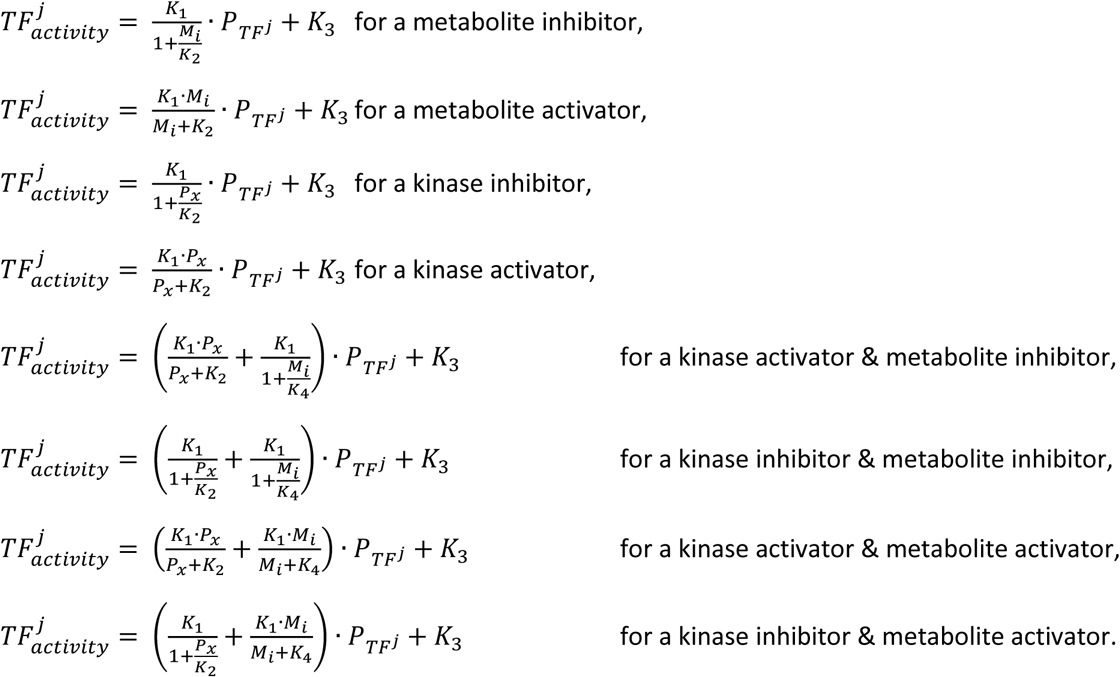

Where K_1,2,3,4_ are free parameters in the model, 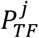 is the relative protein abundance associated to TF^j^, P_x_ represents the relative protein abundance of kinase x and M_i_ the level of metabolite i.

For models in which we assume the non/inhibitory actions of a single metabolite or kinase, we use a non-linear model fitting scheme to find the best set of 3 parameters to describe changes in TF activity across cell lines. To minimize the probability of local minima, we adopted the GlobalSearch function of Matlab and 50,000 starting points. By using the same approach, we tested all possible combinations between pairs of metabolites and kinases that can act either as activators and/or inhibitors, to find the best set of 4 parameters that describe TF activity. For each pair or triplet of TFs and metabolites and/or kinases we estimate the Mean Squared Error (MSE) associated to each tested model. In addition, we estimated the MSE when fixing the model parameters and randomly permuting TF activity, TF protein, metabolite and kinase levels across cell lines (MSE^e^). For each TF we then repeat model fitting, each time by randomly shuffling metabolite and kinases levels. We used this approach to empirically asses the significance of each model in better explaining TF activity. To this end, we calculated MSE values for each model in which kinases and metabolite levels were randomly permuted (MSE_Random_ and MSE^e^_Random_). Finally we retain only those models in which MSE and MSE^e^ are both above the 0.1% of the respective distributions of MSE_Random_ and MSE^e^_Random_. The proteome dataset (ArrayExpress, project accession: E-PROT-2) report protein levels for 100 annotated TFs and 63 kinases, and to reduce computation time we here consider only differentially abundant metabolites annotated to KEGG identifiers. Fitting analysis was performed in Matlab using the “fitnlm” function, on a cluster with 900 computational nodes.

